# Ligation-anchored PCR unveils immune repertoire of TCR-beta from whole blood

**DOI:** 10.1101/007062

**Authors:** Fan Gao, Kai Wang

## Abstract

**Background:** As one of the genetic mechanisms for adaptive immunity, V(D)J recombination generates an enormous repertoire of T-cell receptors (TCRs). With the development of high-throughput sequencing techniques, systematic exploration of V(D)J recombination becomes possible. Multiplex PCR method has been previously developed to assay immune repertoire, however the usage of primer pools has inherent bias in target amplification. In our study, we developed a ligation-anchored PCR method to unbiasedly amplify the repertoire.

**Results:** By utilizing a universal primer paired with a single primer targeting the conserved constant region, we amplified TCR-beta (TRB) variable regions from total RNA extracted from blood. Next-generation sequencing libraries were then prepared for Illumina HiSeq 2500 sequencer, which provided 151 bp read length to cover the entire V(D)J recombination region. We evaluated this approach on blood samples from patients with malignant and benign meningiomas. Mapping of sequencing data showed 64% to 91% of mapped TCRV-containing reads belong to TRB subtype. An increased usage of TRBV29-1 was observed in malignant meningiomas. Also distinct signatures were identified from CDR3 sequence logos, with predominant subset as 42 nt for benign and 45 nt for malignant samples, respectively.

**Conclusions:** In summary, we report an integrative approach to monitor immune repertoire in a systematic manner.

## Background

Activation of the immune system, together with genomic alternation, including somatic hypermutation and recombination, establishes innate and adaptive immunity in the immune system^1,2^. Both the immunoglobulin (Ig) and the T cell receptor (TCR) loci contain many different V, D and J segments, which are subject to a tightly regulated genomic rearrangement process – V(D)J recombination–during early lymphoid differentiation^3–5^. For a given TCR subtype, complementarity determining region 3 (CDR3) is generated by the V(D)J combination at this subtype locus, with the translated protein sequence forming the center of the antigen binding site^6^, defining the affinity and specificity of the receptor for individual peptide-MHC complexes. The CDR3 sequence of dominant clones in cancer patient may serve as a signature to diagnose cancer or to classify tumor into subtypes. In addition, signatures could be obtained at the time of disease diagnosis and then monitored on an ongoing basis, to assess the effects of anticancer therapies or for early detection of recurrence^7,8^

The importance of monitoring TCR in human health and disease has been increasingly recognized. Recent studies showed that TCR repertoire has been found to affect a wide range of diseases, including malignancy, autoimmune disorders and infectious diseases, and, given the broad involvement of the immune system in almost all of human health and disease, this reach should be expected to expand greatly^9.^ However, conventional methods to measure V(D)J recombination have several limitations to prevent detailed characterization of immune repertoire. More recent approaches, such as multiparameter flow cytometry, spectrotyping, or custom-designed real-time PCR assays, are slightly more quantitative and offer higher resolution, but these methods are labor intensive and are unable to offer sequence-level insights as to the exact V(D)J recombination patterns in patients. Solving this problem will enable the wider application of monitoring immune repertoire in clinical settings.

With the development of massively parallel, single-molecule sequencing techniques, it has now become feasible to assay V(D)J recombination by next-generation sequencing, as a means to exhaustively profile the immune repertoire within human subjects. For example, the Roche 454 sequencers has been used to measure and clinically monitor human lymphocyte clonality^10^, which takes advantage of 454’s ability to generate longer sequencing reads that potentially covers V(D)J recombination junction points. Another similar study also used Roche 454 to study human T cell subsets^11^. Note that they separated T cell subpopulations and focused on TCR loci only. However, other investigators have focused on Illumina Genome Analyzer or HiSeq that generates only ∼50 bp to ∼100 bp reads. For example, a group has developed a short reads assembly strategy to first assemble 50bp sequences and then sample the CDR3 diversity in human T lymphocytes from peripheral blood^12,13^. The data analysis involved in such strategy is much less straightforward, but the technology is more accessible and cost-effective in large scale studies. Several additional TCR-Seq studies^13–16^ have been extensively reviewed recently. Some of the commonalities include: (1) they almost always used T-cell DNA or RNA as the first starting material; (2) most of the studies use multiplex PCR reactions to enrich the V(D)J recombinations for next-generation sequencing; (3) large-scale T-cell repertoire analysis has been limited to interrogation of a single TCR subunit per sequencing run, although functional antigen-engaging TCRs are heterodimeric proteins comprising both an α and a ß chain.

In the current study, we explored the feasibility of using ligation-anchored PCR reaction, rather than multiplex PCR reactions, for unbiased amplification of TCR-beta repertoire. The key aspect of our approach is ligating a synthesized linker oligo sequence (/5Phos/NNN…NNN/3ddC/) to the 3’end of single-stranded cDNA, thus it is applicable to extracted RNA samples. As shown below, the method works with frozen whole blood. Additionally, similar to our previous study examining immunoglobulin repertoire^17^, instead of relying on flow cytometry or magnetic beads to isolate T cells or B cell populations from peripheral blood, we attempted to assay RNAs extracted from whole blood directly. Finally, we evaluated sequencing data from the latest Illumina Hi-Seq2500 sequencer, which can generate 151bp single-end reads (∼160 million reads per run), with a significantly shortened turnaround time (48 hours for Rapid Mode) compared to Illumina Hi-Seq2000 or Roche 454 sequencer. Despite the shorter reads than 454 sequencer or Illumina MiSeq sequencer, 151bp read length is still long enough to map V(D)J recombination sites, given appropriate primer design for PCR amplification.

To evaluate this approach, we did a pilot study on patients suffered from meningiomas, the most common primary brain tumors in the United States^18,19^, and compared difference of the detected TRB repertoire between patients with benign (grade I) and malignant (grade III) tumors. This tumor arises from the membranous layers surrounding the central nervous system (CNS), and is not subject to blood-brain barrier. Several previous studies have reported the presence of both humoral^20,21^ and cellular^22,23^ immune responses in patients with meningiomas. Indeed, it has been proposed that frequent antibody response against specific antigens in even benign meningioma can serve as diagnostic targets^20^. Therefore, in addition to testing the technical feasibility, our pilot study has the added value of investigating whether there is increased clonality with increased severity of cancer, and whether different patients may share identical or similar CDR3 sequences in clonal T cells.

## Results

### Total RNA extraction and PCR amplification of TRB variable region

To explore all the actively transcribed immune receptor variable regions within whole blood in an unbiased manner, we developed an integrative approach to extract total RNA from whole blood, followed by a PCR protocol to capture the variable region (V-region) of specific immune receptor (Figure 1A). In this scheme, after total RNA extraction from whole blood, we generate single-stranded cDNA from reverse transcription of mRNA. After ligation of a synthesized linker oligo sequence to the 3’end of single-stranded cDNA, PCR reaction was carried out using a primer pair (primers P-U and P-VDJ) to target the linker sequence and the consensus starting sequence of immune receptor constant region (C-region), respectively. The P-VDJ primer was specifically designed such that it did not have high sequence homology to any other region other than the TCR locus, and that it avoids any known SNPs. In our proof-of-concept study, total RNA extracted from whole blood of a healthy donor was used to test our methodology on capturing TRB variable region (Figure 1B). For samples that were subject to both reverse transcription (either oligo-dT or random primer mix) and linker ligation, PCR reaction resulted in a predominant band with fragment size around 500 bp, roughly the size expected for TRB variable region. For control samples without reverse transcription, this band was not visible. Additionally, controls without linker ligation did not produce this specific band in DNA gel. Thus, the protocol developed in our study should be able to capture transcribed TRB variants from whole blood directly, without separation of T cells first.

**Figure 1.**
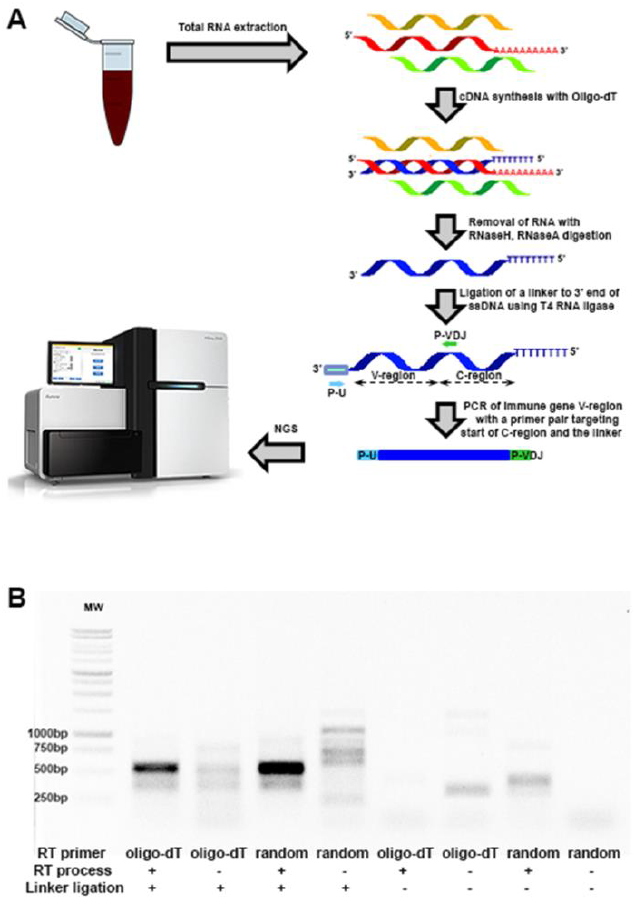
PCR method utilizing single target-specific primer to explore actively transcribed immune genes. A), The scheme of an integrative approach to explore immune repertoire from whole blood (P-U stands for a universal primer sitting on the linker, whereas P-VDJ represents a TRB-specific primer targeting the consensus sequence in the constant region right after J-segment); B), DNA gel image of PCR amplified variable region (V-region) of TRB gene from a healthy donor under different conditions.

### TRB profiling using Illumina high-throughput sequencing

To further evaluate whether the developed protocol is applicable to profile TRB repertoire of clinical samples, we tested the protocol on several blood samples from patients with malignant and benign meningiomas (**Supplementary Tables 1**). The fact that immune responses are present in patients with meningiomas^20,21^ suggests that immune receptors such as TCR may respond to this tumor type and serve as a biomarker for diagnosis. We followed the same protocol to profile TRB V-region variants using previously collected whole blood samples (stored at −80°C for more than one year) from two malignant patients and two benign patients, respectively. Compared to controls without linker ligation, all four reverse transcribed samples with ligation showed the expected predominant band (∼500bp, Figure 2A), suggesting the robustness of this approach. The amplified fragments were further used for next-generation sequencing library preparation.

**Figure 2.**
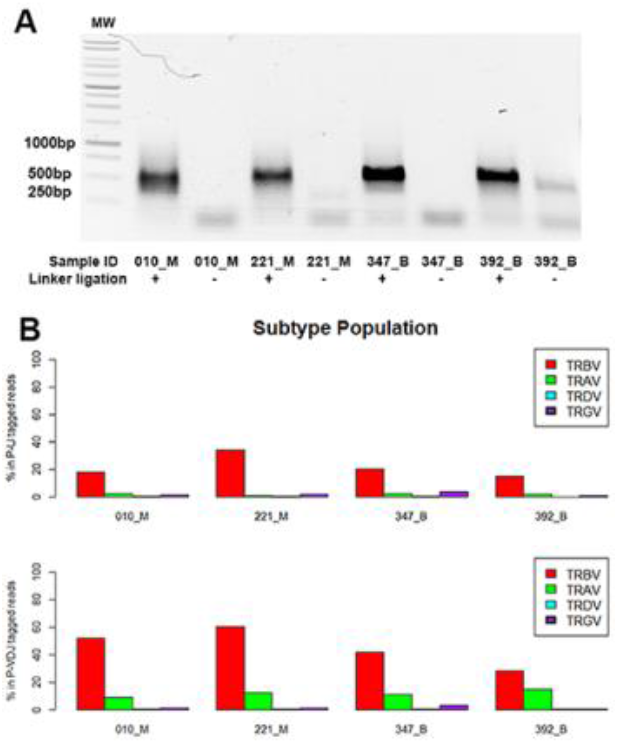
Specificity of the integrative approach in amplifying V-region of TRB gene from whole blood of patients with meningiomas (malignant and benign samples were labeled with M and B, respectively). A), DNA gel image of amplicons from malignant and benign patients under linker ligation or control condition; B), Subtype population of sequenced reads that are mappable to V-region of TCR in each meningioma patient (results are based on the reads with P-U or P-VDJ end tags).

Purified PCR amplicons were barcoded with Illumina TruSeq kit for high-throughput sequencing. The latest Illumina HiSeq-2500 sequencer was used for sequencing, with load of ∼30% PhiX for proper matrixing and phasing into the lane. Collected sequencing data was filtered, and the selected reads were mapped using IgBLAST (see **Methods, Supplementary Tables 2&3**). Of note, the majority of the reads with either P-U or P-VDJ end tag contained TRB fragments (64% to 91% of the mapped TCRV-containing reads belong to TRB subtype). Other subtypes of TCR, such as TRA, TRD, TRG, only accounted for a small portion of the TCRV-containing reads (Figure 2B). Therefore, ligation-anchored PCR approach is indeed capable of profiling TRB repertoire of clinical samples.

### V-segment usage in benign and malignant patients

We next compared the usage of different V-segments from TRB aligning reads (either end-tagged with P-U or P-VDJ primers) in patients with malignant and benign meningiomas. Interesting to note, two malignant samples were grouped together in both P-U and P-VDJ tagged V-segment distribution heat maps, suggesting overall similarity in V-segment usage (Figure 3A). Although most of the V-segments did not present consistent change of percentages in the total population between malignant and benign samples (Figure 3A), one segment TRBV29-1 showed drastic increase in the malignant group (top ranked gene from SAM analysis of MeV software suite, two-class unpaired, FDR<2%), revealing expansion of this particular V-segment in whole blood of malignant patients. Consistent with this discovery, when we further sorted out the translationally productive V(D)J reads within P-VDJ tagged read pool (see Methods), we also noticed that TRBV29-1 level was elevated in malignant samples (Figure 3B), with 5 to 7-fold increase of this population. In addition to TRBV29-1, another V-segment TRBV15 also showed an increase in malignant phenotype. Therefore, the detected clonal expansion of TRBV29-1 possibly reflects human immune response to malignant transformation of meningiomas.

**Figure 3.**
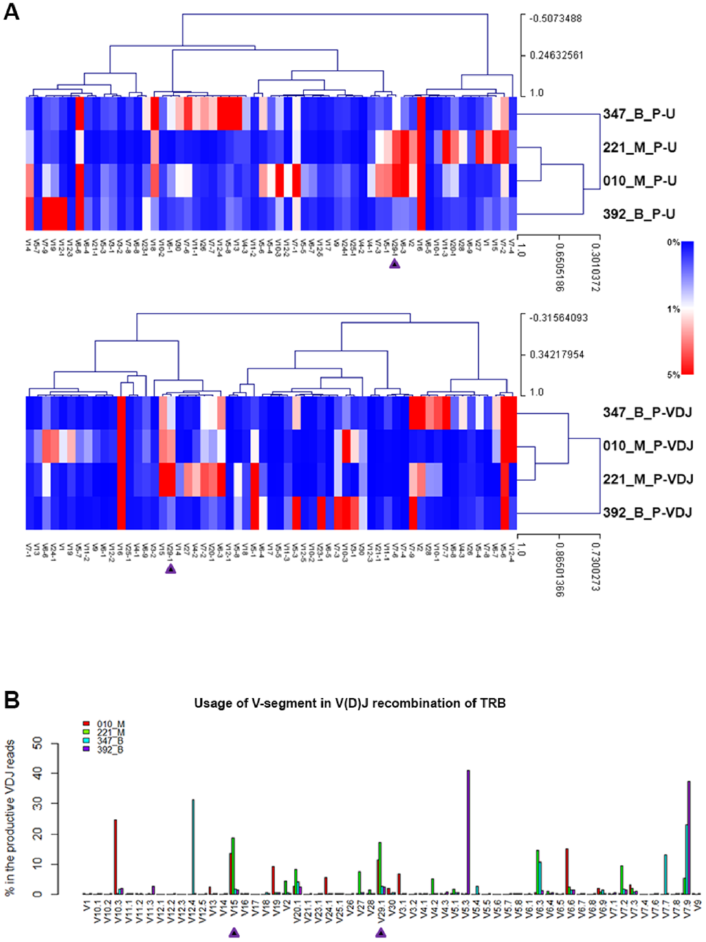
The usage of different V-segments in TRB gene in meningioma blood samples. A), Hierarchical clustering of V-segment usage from both P-U and P-VDJ primer-aligned reads in each patient (color scale represents % of total primer-aligned reads). B), V-segment usage from V(D)J-containing reads that are translationally productive (no stop codon). Arrows highlight expanded V- segments in malignant samples.

### Characterization of “immune signature”–CDR3

In addition to explore potential clonal expansion, we are also interested to evaluate our experimental protocol in capturing the sequences of the CDR3 regions of TCR. From high-throughput sequencing data, we deduced the unique DNA sequence logos of the FR3-CDR3 junction part (Figure 4A) with the last 20 nt of FR3 and the first 60 nt into CDR3 included, and evaluated if any signatures were recurrent features that can separate malignant from benign patients. Initial observation at positions 54 and 56 (Figure 4A) revealed that the most frequent bases were C and G in the malignant samples; however, they switched to base T in the benign samples. Positions 54 to 56 encode an amino acid that is in frame to TRB coding sequence; in malignant samples, the encoded amino acid was Gln (Q), whereas in benign group, it changed to Tyr (Y). We thus generated protein sequence logos of FR3-CDR3 junction (Figure 4B) for further comparison. Of note, a conserved Cys (C) at the end of FR3 region (position 6) was separated from a QXFGXGTRL motif in CDR3 by 11 and 10 amino acids in malignant and benign sample groups, respectively. In a previous study^12^, two major CDR3 subsets with 42 nt and 45 nt were identified in a pooled normal human leukocyte RNA sample. In our study, the protein sequence logos of meningioma blood samples revealed predominant 42-nt subset in benign group and 45-nt subset in malignant group. It is possible that FR3-CDR3 junction presents specific sequence features of different patients with distinct profiles of immune response.

**Figure 4.**
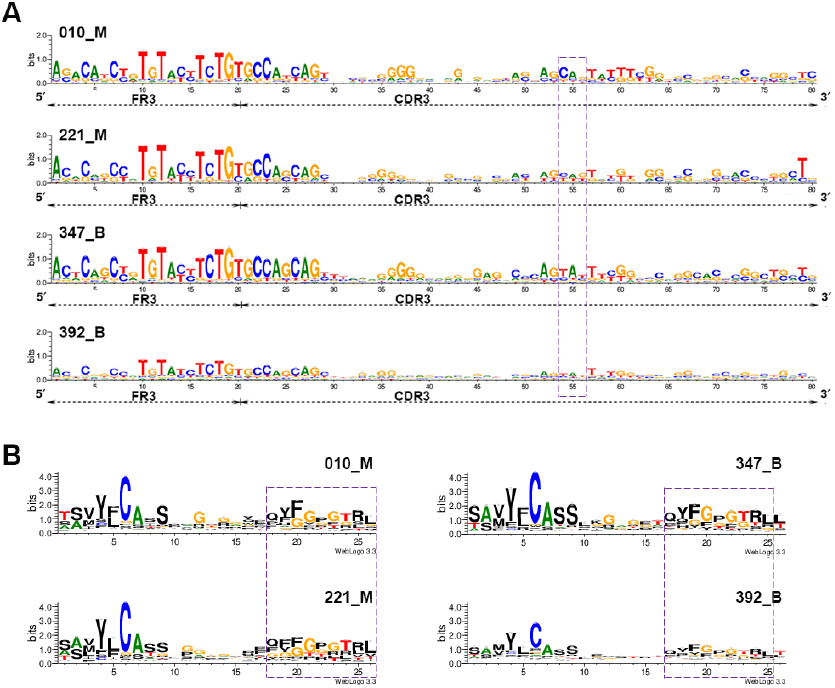
Sequence logos for detected FR3 and CDR3 portions of meningioma blood samples. A) Visualized in the DNA sequence logos at the last 20 nt of FR3 and the first 60 nt into CDR3. A purple-colored box highlights positions 54 to 56 in the DNA sequences. B), In the protein sequence logos, a highly conserved Cys(C) at the end of FR3 is at position 6^th^ of the translated sequence. Purple-colored boxes enclose QXFGXGTRL motif in malignant and benign protein sequences.

### Exploration of V-J segment pairing in V(D)J recombination

To study V-J pairing in V(D)J recombination, normalized pairing frequencies between different V- and J-segments were presented in a Circos plots^24^ (Figure 5). Contrast to only a few predominant V-J pairing events in benign samples, malignant samples showed expansion of more V-J pairing events. Furthermore, pairing of TRBV5-6 with TRBJ1-1 represented one of the major pairing events detected in all samples. Three pairing events including TRBV7-3 with TRBJ2.2, TRBV30 with TRBJ2-6, and TRBV16 with TRBJ1-3, occurred more frequently in the two malignant samples compared to benign ones, possibly representing clonal expansion of specific TCR-expressing T cells in both malignant samples. In summary, despite the use of 151bp sequencing length, our protocol enabled the delineation of the V-J segment pairing in an efficient manner.

**Figure 5.**
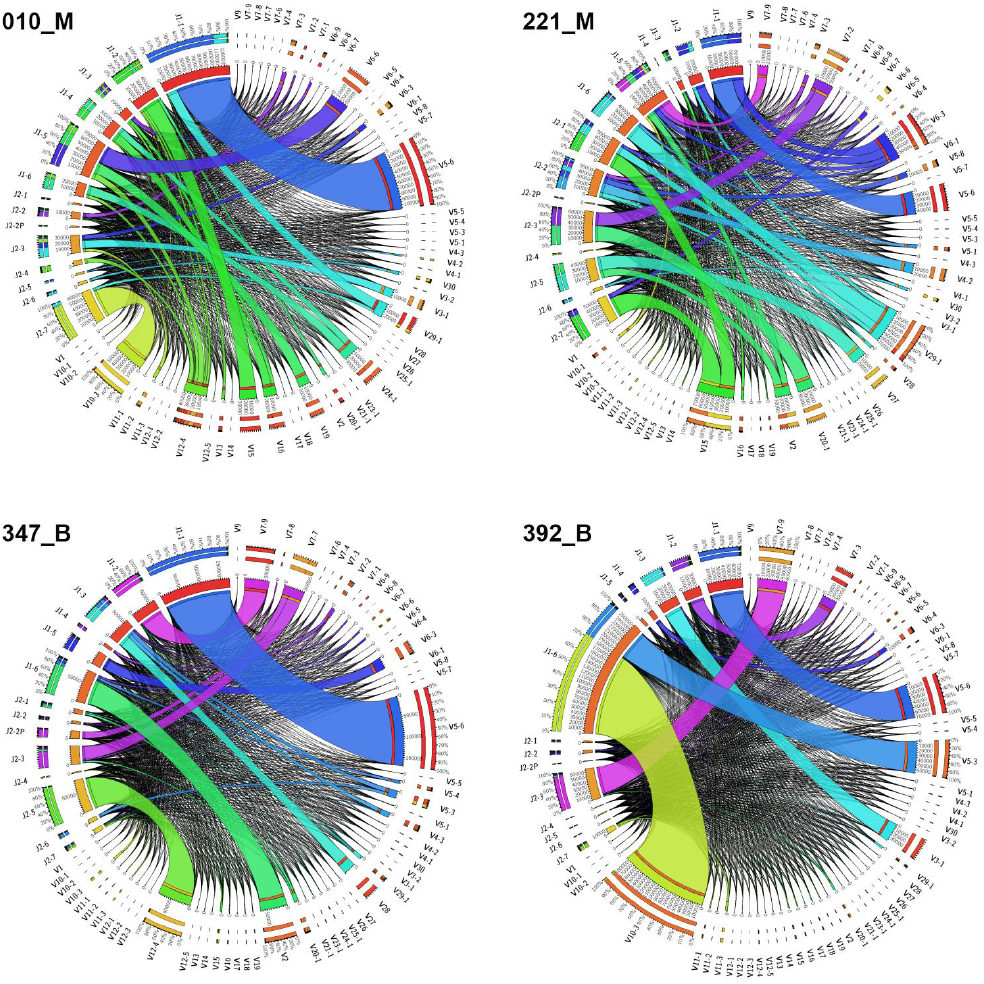
Circos maps illustrating pairing frequencies of V-segments and J-segments from V(D)J-containing reads (all identified segments are ranked by usage frequencies and presented in a clock-wise orientation from the top of the circle, in the order of V-segments and J-segments).

## Discussion and conclusions

In the current study, we presented an integrated approach by using single primer PCR together with next-generation sequencing to interrogate immune repertoire of TCR-beta. We have demonstrated the technical feasibility to use this system to infer immune repertoire, using whole blood from four meningiomas samples. By aligning reads to a sequence database of germline V-genes, D-genes and J-genes, the usage of different V-gene segments was quantified. Interestingly, comparison between malignant and benign groups revealed an increased usage of TRBV29-1 in malignant meningiomas. Analysis of CDR3 region sequence logos revealed addition of 1 amino acid between FR3-CDR3 junction and a consensus motif in malignant group. However, we caution that these observations were made on a small number of samples, and they may not have any biological significance. Our purpose is to use these data to demonstrate the technical feasibility of “single-primer” interrogation of immune repertoire, rather than determining what differs between malignant and benign tumors.

There are several unique aspects of our protocol, compared to previous studies. First of all, total RNA is extracted directly from frozen blood samples for profiling, thus the procedure can be easily adapted for clinical application. Second, by using ligation-anchored PCR for amplification, all the recombination events at a particular immune gene locus can be amplified in an unbiased manner. Finally, sequencing of barcoded libraries through Illumina Hi-Seq 2500 ensures fast turn-around time (less than 48 hours) and good sequencing depth (∼160 million reads per lane) at a relatively low cost.

There are several major limitations of our protocol as well. First, due to the need to add in 3’-adaptors to the cDNA terminus for ligation-anchored PCR, our method relies on RNA samples, which are less readily available and more vulnerable to degradation, compared to genomic DNA samples. However, in the current study, we used frozen whole blood samples and still obtained satisfactory results, suggesting that it is practically feasible to use this method in real-world clinical settings. Second, although Illumina Hi-Seq 2500 provided longer read length (151bp) than Illumina Hi-Seq 2000 (101bp) to cover immune signature region CDR3, longer read length is still needed to cover the entire variable region of immune genes. Thus the latest Illumina MiSeq with 250bp chemistry should be more suitable for profiling immune repertoire. Third, our sequencing data showed higher sequencing depth of malignant sample libraries (010_M, 221_M) compared to benign ones (**Supplementary Table 2**). We acknowledge that uneven mixing of the constructed libraries for sequencing could be the reason. To address the issue of sequencing depth, we performed additional data analysis using the first 7 million sequencing reads of 010_M and 221_M data (**Supplementary Figure, Supplementary Table 4&5**). Of note, the results from the 7-million-read data were consistent with the conclusions from the whole sequencing data, indicating that the sequencing depth in our study is deep enough. Fourth, we specifically designed primers to target TCR-beta isoforms. Our sequencing data contained some other TCR isoforms as well, suggesting the presence of some minor cross reactions. Our future studies may design specific primers for other types of isoforms as well, to investigate whether it is feasible to interrogate all major TCR isoforms in the same sequencing run. Finally, the sample size in our study is small, which prevents us to draw any statistically significant conclusions on clonal expansion of TRB subtypes associated with malignant transformation of meningiomas; however, the results from high-throughput sequencing data analysis do support validity of the developed integrated approach.

In conclusion, we have developed a ligation-anchored PCR approach to interrogate immune repertoire, and demonstrated its technical feasibility and effectiveness in capturing the TCR-beta landscapes. Further development of this technology may enable a comprehensive delineation of immune repertoire, including other forms of TCRs as well as immunoglobulins.

## Methods

### Sample collection, RNA extraction and cDNA synthesis

Peripheral blood from patients with meningiomas from the USC Brain Tumor Bank was collected for the study. For assay optimization purpose, peripheral blood from a healthy donor was also collected. The Institutional Review Board (IRB) at the University of Southern California reviewed and approved the study. Informed consent was obtained for all participants.

Total RNA was extracted from ice thawed 400µL of frozen blood (stored at −80°C) using TRIzol extraction reagent (Life Technologies, Grand Island, NY). For each TRIzol extracted RNA sample, 2mL of anhydrous ethanol was added before passing through an RNeasy-mini column (QIAGEN, Valencia, CA) with in-column DNase I digestion performed. Purified total RNA was eluted in ddH_2_O for further cDNA synthesis using M-MuLV reverse transcriptase kit (NEB, Ipswich, MA) following the recommended protocol, and stored at −20°C.

### Linker ligation and PCR amplification of TRB variants

For each reverse transcribed product, AMPure XP magnetic beads (Beckman Coulter, Danvers, MA) was added to the mixture, followed by 80% ethanol wash to remove primers introduced in reverse transcription step. Each sample was eluted in 30µL EB buffer (QIAGEN, Valencia, CA) with 1µL RNase H (10U/µL, Epicentre, Madison, WI) and 2µL RNase A (5µg/µL, Epicentre, Madison, WI) added to digest RNA templates for 1 hour at 37C, followed by a heat-inactivation step for 5 min. at 95°C.

Ligation of the linker primer to RNase-treated cDNA sample (ssDNA) was carried out in a total volume of 60μL using T4 RNA ligation kit (Promega, Madison, WI) following vendor recommended protocol. The linker primer was synthesized (Integrated DNA Technologies, San Diego, CA) based on the sequence and modification in a previous study^25^. The mixture was incubated overnight at room temperature. For control reaction, ddH_2_O instead of T4 RNA ligase (Promega, Madison, WI) was added.

After overnight incubation, the ligation product was incubated with AMPure XP beads for magnetic purification. Each sample was eluted in 20µL ddH_2_O. PCR reaction was performed using HotStart PCR reagent kit (EMD Chemicals, Billerica, MA) with 2µL of ligated ssDNA as PCR template. In a reaction mixture without primer loaded, a universal primer targeting the linker (GCG GCC GCT TAT TAA CCC) was first added for 2^nd^ strand synthesis at 95°C for 2.5 min., 70°C for 5min. and 4°C forever. Then another primer targeting the consensus sequence of TRB constant region (GAC CTC GGG TGG GAA CAC) was added for 40 cycles of PCR reaction with annealing at 65°C and extension at 70°C.

### NGS library preparation and sequencing

Illumina TruSeq Sample Prep Kit (Illumina, San Diego, CA) was used for library preparation. Briefly, amplicons were purified before sequential steps of end-repair, adenylation of 3’end and adaptor ligation using the recommended protocol. Finally, the barcoded libraries were enriched with 10 cycles of PCR reactions and cleaned with AMPure XP beads. The quality of prepared libraries was assessed using Agilent High Sensitivity DNA Chip on a Bioanalyzer (Agilent Technologies, Santa Clara, CA). The bar-coded libraries were quantified with KAPA Library Quantification Kit (Kapa Biosystems, Woburn, MA) before mixing for sequencing in a single lane on the Illumina HiSeq 2500 system. The raw FASTQ data files of 151 bp single-end reads were collected for downstream analysis.

### Analysis of sequencing data

For the raw FASTQ data collected from high-throughput sequencing, sequencing quality analysis was first performed using FASTQC^26^ to ensure the mean values of sequence quality (Phred Score) for each base is greater than 32. Then a custom PERL script was used to select the reads with the starting sequences containing either P-U (with additional 5 bp extended to the ligation site) or P-VDJ primer sequence. Unlike genome or exome sequencing, the nature of sequencing data from immune gene variable regions requires 1) mapping against a sequence database containing V-gene, D-gene and J-gene segments; and 2) aligning the V(D)J junctions formed. For mapping of the sequenced reads, recently released IgBLAST^27^ was employed to identify reads that can be mapped to germline V, V-D, V-J, or V-D-J segments. The default parameters were used for mapping. Due to lack of a standalone version of IgBLAST for TCR gene analysis, the sequencing data (reformatted to FASTA files) were split into small batches and uploaded to the IgBLAST web server for mapping. Hierarchical clustering analysis was carried out using MEV, which is a component of the TM4 suite of microarray analysis tools^28^. Sequence logo graphs were generated using standalone version of WebLogo (weblogo.berkeley.edu)^29^. Circos maps were plotted using Circos Table Viewer web server (http://mkweb.bcgsc.ca/tableviewer/)^24^. All other plots were generated using R.

## Acknowledgements

We are grateful for the patients with meningiomas for donating their blood samples for research purposes. We thank William J. Mack for help in obtaining whole blood samples and phenotype information.

## Authors’ contribution

FG & KW designed the study. FG performed the experiment, collected and analyzed the data. FG & KW wrote the paper. All authors read and approved the manuscript.

